# Temporal environmental variation imposes differential selection on genomic and ecological traits of virtual plant communities

**DOI:** 10.1101/2020.03.24.005058

**Authors:** Ludwig Leidinger, Juliano Sarmento Cabral

## Abstract

The reaction of species to changing conditions determines how community composition will change functionally — not only by (temporal) species turnover, but also by trait shifts within species. For the latter, selection from standing variation has been suggested to be more efficient than acquiring new mutations. Yet, studies on community trait composition and trait selection largely focus on phenotypic variation in ecological traits, whereas the underlying genomic traits remain relatively understudied despite evidence of their role to standing variation. Using a genome-explicit, niche- and individual-based model, we address the potential interactions between genomic and ecological traits shaping communities under an environmental selective forcing, namely temporal variation. In this model, all ecological traits are explicitly coded by the genome. For our experiments, we initialized 90 replicate communities, each with ca. 350 initial species, characterized by random genomic and ecological trait combinations, on a 2D spatially-explicit landscape with two orthogonal gradients (temperature and resource use). We exposed each community to two contrasting scenarios: without (i.e. static environments) and with temporal variation. We then analyzed emerging compositions of both genomic and ecological traits at the community, population and genomic levels. Communities in variable environments were species poorer than in static environments, populations more abundant and genomes had a higher numbers of genes. The surviving genomes (i.e. those selected by variable environments) coded for enhanced environmental tolerance and smaller biomass, which resulted in faster life cycles and thus also in increased potential for evolutionary rescue. Even under the constant environmental filtering presented by temporal environmental variation, larger, more linked genomes allowed selection of increased variation in dispersal abilities. Our results provide clues to how sexually-reproducing diploid plant communities might react to increased environmental variation and highlights the importance of genomic traits and their interaction with ecological traits for eco-evolutionary responses to changing climates.

## Introduction

Communities of plant species are the result of different abiotic and biotic conditions (Huntley 1991). Changes in those conditions will therefore also reflect on communities and their trait composition. Response strategies that enable species survival under changing conditions may vary across species. They can, for instance, select for survival (Holt 1990), for lower body mass (Parmesan 2006), for dispersal (Berg et al. 2010), or for adaptation to new conditions (Joshi et al. 2001, Jump and Peñuelas 2005, Bell and Gonzalez 2009). Given enough time, this will result in the communities passing through ecological species successions (Huston and Smith 1987) and evolutionary taxon cycles (Ricklefs and Bermingham 2002). Even in short periods, populations within communities can change their traits in response to environmental variation via rapid evolution (Maron et al. 2004). In this case, selection on standing variation can be more efficient than aquiring novel mutations (Barrett and Schluter 2008, Bolnick et al. 2011). This standing variation can be both intraspecific and intra-individual, i.e., within-genome variation. A high standing variation thus provides a resource for populations to quickly respond to changing environments (Cochrane et al. 2015). However, the genomic traits which enable and maintain standing variation remain largely understudied in ecological and eco-evolutionary studies (but see Schiffers et al. 2012, Matuszewski et al. 2015).

Many functional species traits are quantitative and subject to genetic interactions, such as epistasis, pleiotropy and genetic linkage. To infer a direct connection between phenotype and genotype is therefore complex (Korte and Farlow 2013). Still, all this genomic background determines standing genetic variation, which in turn will constrain which individual phenotypes are possible and thus a population’s evolutionary potential. With the increasing availability of exhaustive genetic data, considering detailed genetic factors in eco-evolutionary models has become more feasible, especially for model species (Frachon et al. 2019, Exposito-Alonso et al. 2019). Indeed, there is an increasing amount of genetic data at the population or even at the individual level (e.g. Domingues et al. 2012, Alonso-Blanco et al. 2016). Nevertheless, manipulating real-world systems to conduct meaningful experiments to isolate factors on both functional and genetic levels is difficult (but see Booth and Grime 2003). Thus, although the importance of genetic factors for ecological processes has long been recognised (Holt 1990), investigating its effects in real-world systems remains challenging (Hughes et al. 2008).

Simulation models provide a powerful alternative to overcome the empirical challenges of investigating and manipulating genetic traits and all the trait-mediated ecological functions they control. Modeling studies can cover any organisational level in biology, from genomes over species to communities (Matuszewski et al. 2015, Kubisch et al. 2014, Münkemüller et al. 2012, Saupe et al. 2019), and thus are suitable tools to explore potential eco-evolutionary regulations of species traits. Therefore, we developed a Genome-explicit Meta-community Model (GeMM, Fig. 1) to address the interplay of genomic and ecological traits in species communities under an environmental selective force, namely temporal environmental variation. Specifically, we address the following questions. (a) Which ecological and genomic traits enable survival in temporally variable environments? (b) How do temporally variable environments shape standing variation (phenotypic and genetic)? We designed a simulation experiment under two different environmental scenarios, namely with versus without temporal environmental variation (variable and static environments, respectively) and analyzed genomic and ecological trait characteristics of surviving communities. We expected communities in variable environments to select for higher tolerances (Holt 1990), higher dispersal abilites (Berg et al. 2010) and lower biomass (Parmesan 2006) and to exhibit increased standing variation, both genetic and phenotypic (Cochrane et al. 2015). While our expectations on trait responses were largely confirmed, we find that standing variation is decreased for most traits except dispersal. Our findings on virtual communities suggest how eco-evolutionary dynamics of real plant communities might unfold under changing environments.

**Figure 1.**
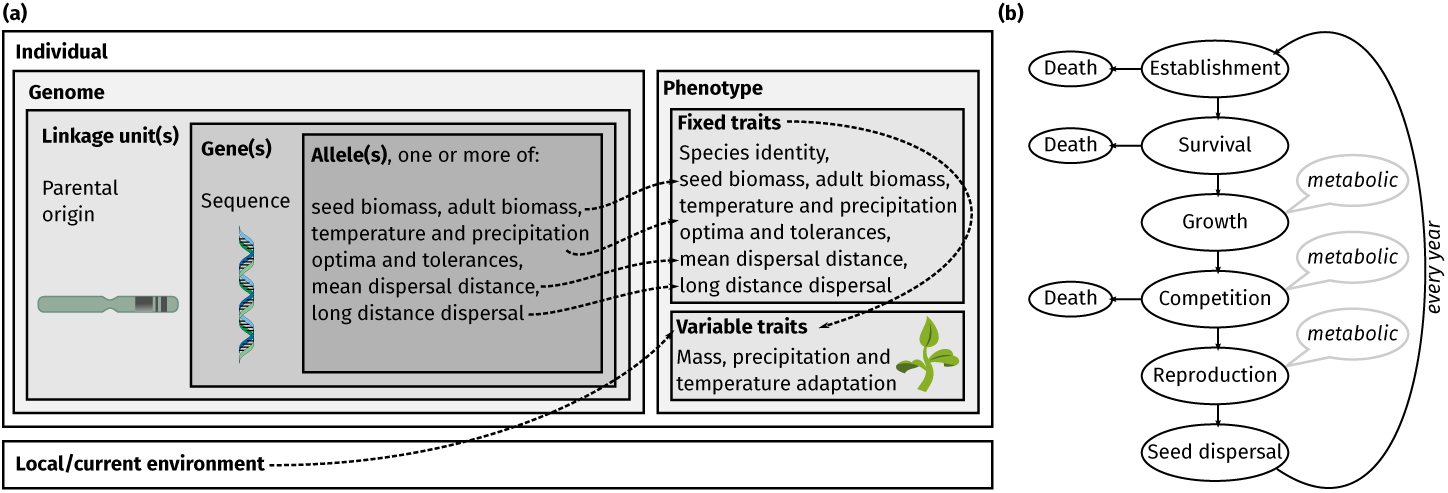
Schematic of the model. (a) Individuals represent the base agents in the model. They are comprised of a phenotype which interacts with other individuals and the environment, and a genome. The genome is diploid and consists of maternal and paternal sets of linkage units, which combine genes as one hereditary unit. Each gene may code for one or more alleles of functional traits. The expressed trait in the phenotype results as the average of all associated alleles in the genome. The expression of some of the traits (“variable traits”) additionally depends on the local current environment and may change over time. (b) Flow of processes each individual passes every year. Some of the processes are dependent on the local temperature and individual biomass (marked “metabolic”), while all processes depend on an individual’s phenotypic traits (see (a)). Dashed arrows represent influences, solid arrows represent sequence of events.

## Materials and Methods

### The model

#### General structure

We use GeMM (version 1.0.0) — a genome- and spatially-explicit, niche- and individual-based model for plant metacommunities written in julia (Bezanson et al. 2017, Fig. 1). The model generates metacommunity dynamics (Hanski 2001, Leibold et al. 2004) and it considers explicit local population and community assembly dynamics emerging from genomic and individual level processes. The model simulates discrete time steps, which can be translated to one year. In the model, individuals belong to species, which are characterized by individuals with identical genetic architecture (i.e. genome size and linkage) and ecological traits (dispersal ability, environmental niche and size) falling within a species-specific Gaussian trait distributions (Fig. 1 (a)). Thus, individuals of the same species are not functionally identical, depicting intraspecific phenotypic variation. Dispersal of individuals (i.e. seeds) interconnects grid cells, while the position of individuals is characterized by the grid cell coordinates, i.e., all individuals are concentrated in the center of the respective grid cell.

#### Eco-evolutionary processes

Like some previous ecosystem models (Harfoot et al. 2014, Cabral et al. 2019, see Cabral et al. 2017 for a review), yearly vegetative growth in biomass, fertility and mortality rates in the model are controlled following the metabolic theory of ecology (MTE, Brown et al. (2004), Price et al. (2010)). Accordingly, the model considers discrete yearly time steps. In MTE, a biological rate *b* depends on the temperature *T* and individual mass *M*, scaling a base rate *b*_0_ as:

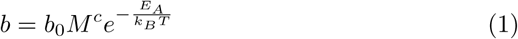

where *E*_*A*_ is the activation energy and *k*_*B*_ the Boltzman constant. The exponent *c* is 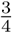 for biomass growth and reproduction, and −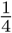 for mortality (Brown et al. 2004). This results in smaller individuals having a higher mortality than bigger ones, while individuals in cooler conditions have a lower mortality than those in warmer conditions. Using the MTE means reduced parameterization effort, since *b*_0_ values for the different processes are global constants and thus identical for every species. Additionally, the emerging longevity-fecundity trade-off that comes with mass-regulated rates has been shown to inherently supress the evolution of “super-species” (Cabral et al. 2019).

Over the course of a simulation, individuals thus grow in size, passing three life stages: (1) seed, (2) juvenile, and (3) adult. Individuals disperse as seeds, establish, grow and become reproductive adults (Fig. 1 (b)). Both seed biomass and adult biomass, i.e., the threshold biomass where individuals become reproductive, are two of the central, genetically-coded traits that define individuals (Fig. 1 (a), Table 1). Adults are monoaecious and reproduce sexually with a random adult of the same species whithin the same grid cell to produce new seeds. Seed dispersal follows a logistic dispersal kernel with genetically-coded mean dispersal distance and shape parameter *µ* and *s*, respectively (see Bullock et al. 2017). In our discrete landscapes, dispersal is modeled as centroid-to-area, with expected mean dispersal distances usually around equal to the length of the grid cells (cf. Chipperfield et al. 2011). Furthermore, all individuals have encoded preferences concerning two different environmental measures: the first, temperature, has a direct effect on biological rates, as described by the MTE (Brown et al. 2004) and affects density-independent mortality, while the second is a surrogate for environmental resources, e.g., water. Thus, from here on this second axis is called precipitation for simplicity. Individuals’ adaptation to precipitation conditions determine their competitive abilities. Both these preferences are characterized by an optimum and a tolerance, which are represented as mean and standard deviation of a Gauss curve, respectively. The degree of mismatch between an individual’s preference optimum with the local environment (i.e. within the grid cell) determines its adaptation value (i.e. environmental fitness). Near their optimum, individuals with higher niche tolerance have lower adaptation values than individuals with narrower breadth (i.e. specialists, Griffith and Sultan (2012)). During establishment, the adaptation values toward temperature and precipitation are calculated for each new seed based on the local conditions and phenotypic traits (Fig. 1 (b)). Furthermore, each time environmental conditions change, all individuals in the affected grid cell pass establishment again to re-calculate their adaption values. These adaptation values are functional for two different subsequent processes. First, individuals experience a metabolic, density-independent mortality (Brown et al. 2004). This mortality further scales with individual temperature adaptation, so that mortality is higher for individuals which are poorly adapted to the surrounding temperature (Cook 1979). Second, all individuals in a cell compete for the limited available space in the grid cell, i.e., total sustainable biomass. If the combined biomass of all individuals in a cell exceeds the grid cell’s carrying capacity biomass, individuals are removed from the community until biomass is within grid cell limits. The choice which individuals to remove is based on pair-wise comparisons of random pairs of individuals. From any of such two individuals, the individual less adapted to local precipitation conditions is removed.

**Table 1.** Model parameters, their meaning and relevance. Phenotypic traits *y* 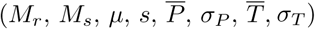 are always the average of all corresponding trait loci *y*_*l*_ in the genome. Values are arbitrary, but within empirically or theoretically supported ranges (see main text and supplementary materials for details) and dimensionless unless otherwise specified. The variability column describes whether and how values might change. Constant: values are global constants across scenarios; genome: values might differ within an individual’s genome, potentially giving rise to different phenotypes; scenario: values differ between scenarios, but stay constant within scenarios; species: values might differ between species, but stay fixed within species. SD: standard deviation

| Parameter | Description | Value or Range | Variability |
| --- | --- | --- | --- |
| $E_A$ | Activation energy | $1 \times 10^{-19}$ J (adapted from Brown et al. 2004) | constant |
| $k_B$ | Boltzman constant | $1.38 \times 10^{-23}$ J K<sup>-1</sup> | constant |
| $K$ | Carrying capacity | 100 kg | constant |
| $r_0$ | Base fecundity | $1.4 \times 10^{12}$ (modified after Brown et al. 2004) | constant |
| $g_0$ | Base growth rate | $8.8 \times 10^{10}$ (modified after Brown et al. 2004) | constant |
| $m_0$ | Base mortality rate | $1.3 \times 10^9$ (modified after Brown et al. 2004) | constant |
| $\delta_P$ | Temporal precipitation SD | 0.0 or 0.2 | scenario |
| $\delta_T$ | Temporal temperature SD | 0.0 °C or 0.2 °C | scenario |
| $n_l$ | Number of loci | 1 to 20 (Fournier-Level et al. 2011) | species |
| $n_u$ | Number of linkage units | 1 to $n_l$ | species |
| $\sigma_l$ | SD among trait loci | 0 to $0.1 \times \text{mean of trait}$ | genome |
| $M_r$ | Biomass at reproductive stage | $e^3$ g to $e^{14}$ g (Brown et al. 2004) | genome |
| $M_s$ | Biomass at seed stage | $e^{-2}$ g to $e^{10}$ g | genome |
| $\mu$ | Dispersal kernel mean | 0 to 1 grid cells | genome |
| $s$ | Dispersal kernel shape | 0 to 1 grid cells | genome |
| $\bar{P}$ | Precipitation optimum | 0 to 10 | genome |
| $\sigma_P$ | Precipitation tolerance | 0 to 1 | genome |
| $\bar{T}$ | Temperature optimum | 10 °C to 40 °C | genome |
| $\sigma_T$ | Temperature tolerance | 0 °C to 1 °C | genome |

#### Genetic architecture

All of the aforementioned traits (see Table 1) are coded by one or more genes in an individual’s diploid genome (polygenes). Single genes can also be associated to several traits at the same time (pleiotropy, Solovieff et al. (2013)). Thus, each trait can be represented more than once in the genome (i.e. through different genes at different loci). Since trait representations are subject to species-specific variation, they can constitute different alleles — both within the haploid genome at different loci, but also between the maternal and paternal haploid genomes or between individuals (cf. Nevo 1978). Realized ecological traits *y*, i.e., an individual’s phenotype, are then determined quantitatively by considering all respective loci *y*_*l*_ within an individual’s genome and taking their average. This results in a random degree of species-specific phenotypic and genetic, i.e., intra-individual or intra-genomic, trait variation (cf. Mackay 2001). Lastly, genes may be combined to form a linkage unit, which represent a set of spatially close genes within the same chromosome arm. Linkage units thus comprise the smallest hereditary entities (Hermann et al. 2013, Lande 1984). Haploid gametes receive a complete random set of those linkage units following a recombination process, where each linkage unit can originate from either the paternal or maternal chromosomal complement of the individual producing the gamete. During reproduction, the gametes of two mating individuals thus form an offspring’s (i.e. seed) genome. The phenotypic characteristics of each offspring are then calculated on the basis of its recombined genome and local environmental conditions (Fig. 1 (a)).

A detailed model description with justification for assumptions, equations and parameter values can be found in Supplementary material Appendix 1 (Grimm et al. 2006, 2010). Model parameters are summarized in Table 1.

### Experimental design

#### Simulation arena

We set our simulation experiments in a rectangular landscape of a grid of 5 by 7 grid cells (Fig. 1). Each grid cell had a carrying capacity of 100 kg of total biomass, which approximately relates to 100 m^2^ of grassland (Deshmukh 1984, Bernhardt-Römermann et al. 2011). Landscape size and carrying capacity was arbitrary but ensured computational feasibility. Two perpendicular environmental gradients (temperature and precipitation) ran along the long and short axis of the landscape, respectively. The rectangular shape of our simulation arena provided a longer gradient in the physiologically important temperature direction.

#### Initialization

We initialised each grid cell of the landscape with a different local community of random species. The species characteristics (i.e. genomic and ecological traits) as well as local abundances were chosen randomly from large ranges of uniform-distributed values. On the genomic level, species differed by the number of loci, *n*_*l*_ (maximum = 20, cf. Fournier-Level et al. 2011, Schiffers et al. 2012), intragenomic variation between trait values, i.e., genetic variation, *σ*_*l*_ (maximum = 0.1×trait value), and number of linkage units, *n*_*u*_ (between one and *n*_*l*_, Table 1). To obtain the ecological characteristics of a species, first an average phenotype was defined by randomly selecting a value for each phenotypic trait. These traits, more specifically, the adult biomass trait, were then used to calculate the number of offspring a single individual of this species would have. Given an already determined genetic architecture (i.e. *n*_*l*_, *n*_*u*_, and *σ*_*l*_), each individual of a species was then initialized as follows. For each trait representation (i.e. gene) within the genome, the associated trait value was chosen randomly following a Normal distribution with the trait value of the average phenotype as mean and standard deviation the product of *σ*_*l*_ and the trait value (Table 1). Afterwards, the initial phenotype for each individual was calculated based on all genes in the genome. This resulted in varying degrees of intragenomic and intraspecific standing variation. We disabled mutations in our experimental design so that this standing variation was the only resource for selection. Grid cells were then filled with populations of several species until carrying capacity was reached. Because species vary randomly in their traits, including biomass, initial grid cell communities varied in richness. This resulted in initial communities with on average 10 species per grid cell and a total of 350 species in the landscape.

Values for simulation, global and species-specific parameters that were not varied in the different experimental scenarios were chosen to ensure plausible patterns, most importantly to achieve species co-existence by adjusting the mortality-to-fecundity ratio. Species-specific parameter values were drawn at random from a range that extended beyond what would be realisable in simulations to reduce geometric artifacts within the parameter space (Table 1). This also kept the need for additional assumptions at a minimum, since viable species emerged via environmental filtering and ecological interactions. Global parameter values were either adapted from the literature (Brown et al. 2004, Fournier-Level et al. 2011) or fine-tuned via trying out a range of realistic values.

#### Scenarios

For investigating our general study question about the interplay of environmental variation and ecological and genomic traits, we designed two scenarios. In the first, temperature and precipitation gradients arbitrarily ranged through constant values of 16.85 °C to 22.85 °C (290 K to 296 K) and 3 to 7 (arbitrary quantity), respectively, during the entire simulation run (“static environment”). In the second, initial temperature and precipitation values were the same as in static environments, but could change at each year (“variable environment”). The change followed a gaussian random-walk trajectory to yield positive auto-correlation (Fung et al. 2018). The amount of change (*δ*_*P*_ and *δ*_*T*_, Table 1) was drawn randomly from a Normal distribution with a standard deviation of 0.2. This value corresponds to a moderate rate of change of no more than 0.5 degrees per year in the majority (ca. 99 %) of cases, which we found by trying different values to produce noteable environmental change that did not kill all individuals in a short amount of time. Since our simulation arena represents a small spatial scale, all grid cells changed always by the same value at each timestep. The change of temperature was independent from that of precipitation and vice-versa. Confounding effects, such as landscape configuration, different temporal dynamics, complex dispersal behavior and macro-evolutionary processes (e.g. clade diversification) have been studied elsewhere and were thus not included in the present study (Münkemüller et al. 2012, Kubisch et al. 2014, Aguilée et al. 2018). Table 1 contains the parameters which were varied for the scenarios, their meaning and their values. We simulated 90 different replicates. Each replicate terminated after 1000 simulated years. This duration was adequate to allow quasi-equilibrium (see Results) and short enough to warrant our selection-on-standing-variation rationale (Hermisson and Pennings 2005). Each replicate, i.e., each unique initial community, was subjected to both scenarios. This yielded 180 simulations in total.

We recorded the complete state of the individuals in our simulation world at the start and every 50 years of a simulation run. This data encompassed individual phenotypic and genotypic values. Thus, for every year, we tracked the state of local species populations including location, abundance, demographics, median adaptation, and trait values for all ecological and genomic traits.

### Analyses

To make the individual information more accessible, we calculated summary statistics at the population level. We defined a population as a group of conspecific individuals co-occurring in the same grid cell. For each population, we then calculated median values of each phenotypic trait, the variance of each phenotypic trait (phenotypic intraspecific variation), and medians of the individual genetic variation in each trait. We scaled all variance values by the respective population-specific medians to get coefficients of variation of the median (CV median). In order to compare emerging ecological patterns and identify when equilibrium is reached, we calculated a set of ecological metrics, namely species-richness, i.e., the average number of species per grid cell, *α* (*α*-diversity), the total number of species across the landscape, *S* (*γ*-diversity), *β*-diversity, *β* = *S/α* −1 (Whittaker 1960), population demographic structure (i.e. number of juveniles and number of adults) and range-filling from the data on surviving communities. For diversity indices, we converted our data to community matrices and analyzed them using vegan (Oksanen et al. 2018) in R (R Core Team 2019). To assess demographic structure within communities, we analyzed average numbers of juveniles and adults. Range-filling was calculated as the fraction of grid cells that was occupied by a species over all the grid cells that were potentially suitable for the given species. Suitability was asserted as an arbitrary cut-off where environmental parameters (temperature and precipitation) fell within a species’ tolerance (optimum ± tolerance).

For our study questions, we analyzed the trait composition of surviving communities genomic trait composition (study question (a)), and differences in phenotypic and genetic standing variation (study question (b)) between environments. Since we were interested in general patterns of the effect of environmental variability, rather than the effects of warming or cooling trends, we excluded precipitation and temperature optimum traits from our analyses. We transformed trait and variation distributions before analysis and visualization using a log (*x* + 1) transformation, because they were strongly left-skewed and contained values *<* 1. Additionally, we calculated the degree of genetic linkage as 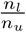, because due to our method of initializing species, *n*_*u*_ directly depended on *n*_*l*_.

To ascertain whether and how trait composition differs between environmental conditions (study question (a)), we first compared species numbers and identities. Because each community is subjected to both environments, we analyzed what proportion of species was shared by both environments, and which were unique to one of the environments. To assess how ecological and genomic traits respond to variable environments, we compared trait characteristics between scenarios by performing principal component analyses on the population trait data. This way, we were able to describe general patterns in trait space between scenarios by relating the total trait space shift to the principal components and correlated trait axes. Additionally, we compared community trait distributions pairwise between environments to identify trends in traits specific to the environments. For this, we calculated linear mixed models using the R package lme4 (Bates et al. 2015) with trait as response, environment as fixed effect and replicate as random effect.

To find out whether there is a selective force on standing variation (both phenotypic and genetic) specific to environmental conditions (study question (b)), we compared the phenotypic and genomic trait variances of surviving communities between scenarios for all traits in separate. We again calculated linear mixed models, with trait variances as response, environment as fixed effect and replicate as random effect.

The model code, experiment definition files and analysis scripts are available at https://github.com/lleiding/gemm. Albeit reporting of significance values is generally inappropriate for simulation models (White et al. 2014), we use significance here to identify which responses are stronger than others.

## Results

### Differences of ecological patterns between environments

Surviving communities in our simulation experiments (Fig. 1) differed in a number of ecological characteristics. Compared to communities in static environments, communities in variable environments were only about half as species-rich on a local level (*α*-diversity, Fig. 2(a)) and exhibited less *β*-diversity (Fig. 2(b)), which resulted in decreased species richness on a regional scale (*γ*-diversity, Fig. 2(c)). Summing over all replicates, a total of 108 species survived in both enviroments, while 256 and 64 surviving species were unique to static and variable environments, respectively. Emerging functional differences comprised higher total abundances in all demographic stages (Fig. 2(d), (e)) and decreased range filling for communities in variable environments (Fig. 2(f)). While all aforementioned metrics were constantly changing during the entire simulation course in the variable environments, in static environments they reached a quasi-equilibrium by year 500. For the following trait-based analyses, we thus used data from that year.

**Figure 2.**
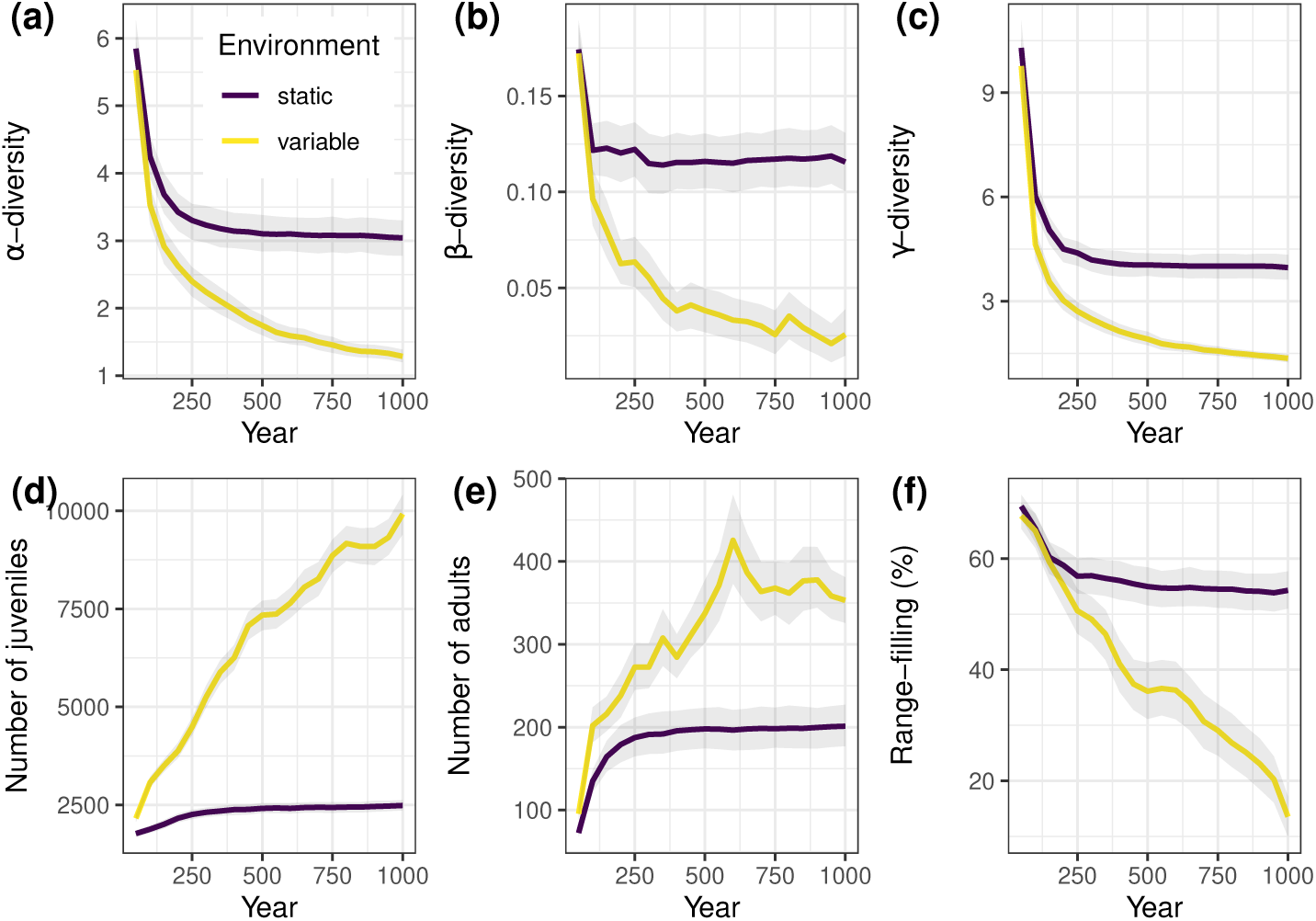
Averaged ecological patterns across the entire simulation arena over time after initialisation. Dark/violet: static environment, light/yellow: variable environment. Grey ribbons represent 95% confidence intervals. (a) local species richness (*α*-diversity) as numbers of species, (b) *β*-diversity (Whittaker 1960), (c) total species richness (*γ*-diversity) as numbers of species, (d) mean number of juveniles, (e) mean number of adults, (f) range-filling, i.e., the percentage of potentially suitable habitat that is actually occupied. Spikes are due to single replicates with extreme values.

### Response of ecological and genomic traits

Surviving communities showed subtle differences in their trait syndromes combining all traits in a PCA. In the first two principal components, populations from variable environments occupied for the most part a subset of the trait space of populations from static environments (mostly overlapping ellipses in Fig. 3). Nevertheless, the trait space of variable environment communities was shifted towards increased environmental tolerances and dispersal abilities and decreased mean genetic variation (negative direction of second principal component - Fig. 3). With the exception of the first, all principal components and thus correlated traits, contributed similarly to the overall explained variance (Supplementary material Appendix 1 Fig. A2).

**Figure 3.**
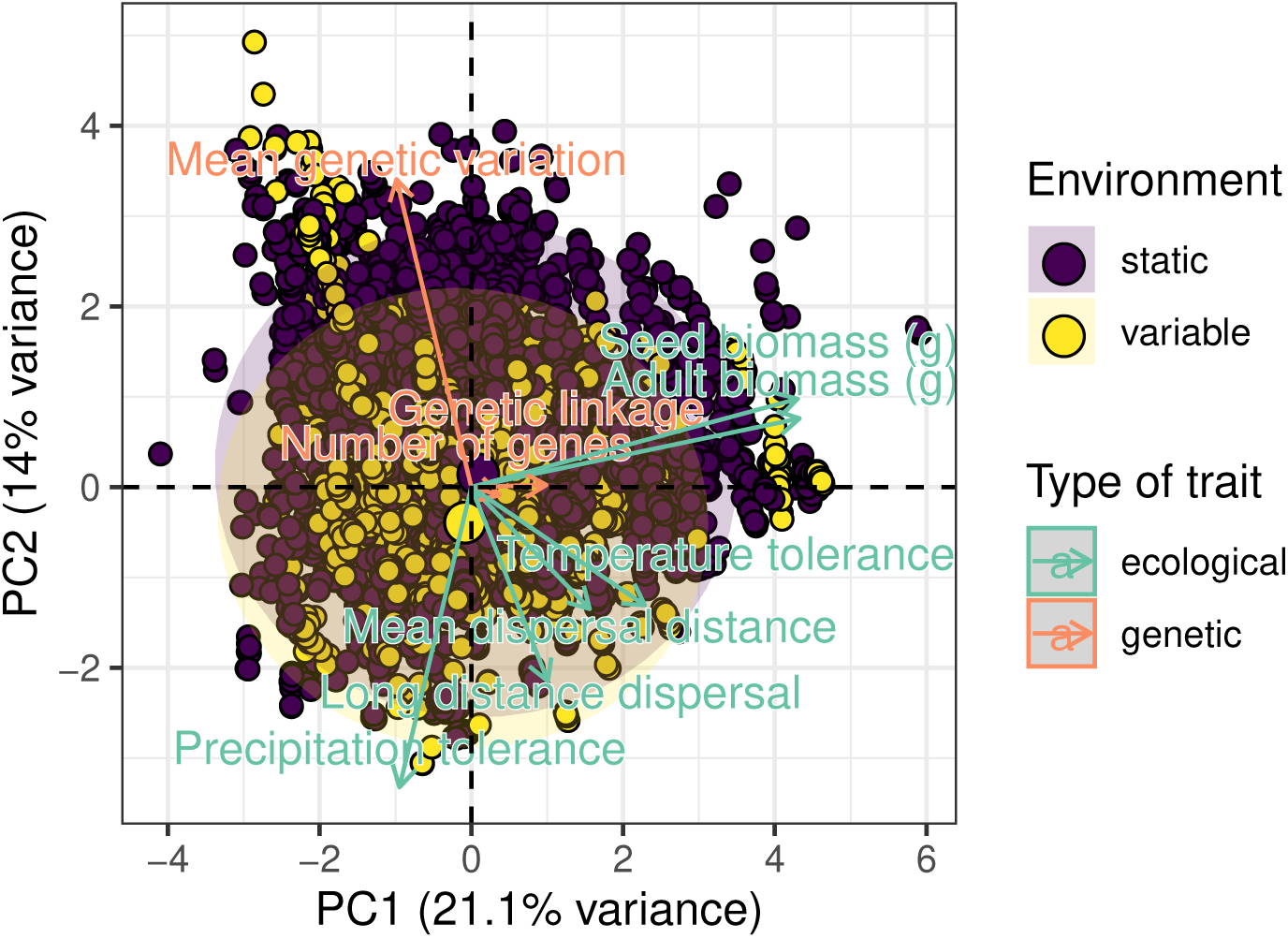
Principal component analysis (PCA) showing trait space characteristics (ecological and genomic) of surviving populations. Biplot of surviving populations and trait axes along the first two principal components. Populations without temporal environmental variation (dark/violet) vs. with temporal environmental variation (light/yellow). Shadowed ellipses highlight areas of 95 % confidence.

Focusing on single traits, communities showed several differences between the two types of environments (Fig. 4(a), Supplementary material Appendix 1 Table A3). Compared to static environments, surviving communities in variable environments showed on average an increased number of genes (*n*_*l*_), increased precipitation and temperature tolerances (*σ*_*P*_ and *σ*_*T*_, respectively), increased long distance dispersal (*s*), decreased adult biomass (*M*_*r*_), and decreased genetic variation (*σ*_*l*_). Seed biomass, mean genetic variation and genetic linkage exhibited no significant differences (Supplementary material Appendix 1 Table A3).

**Figure 4.**
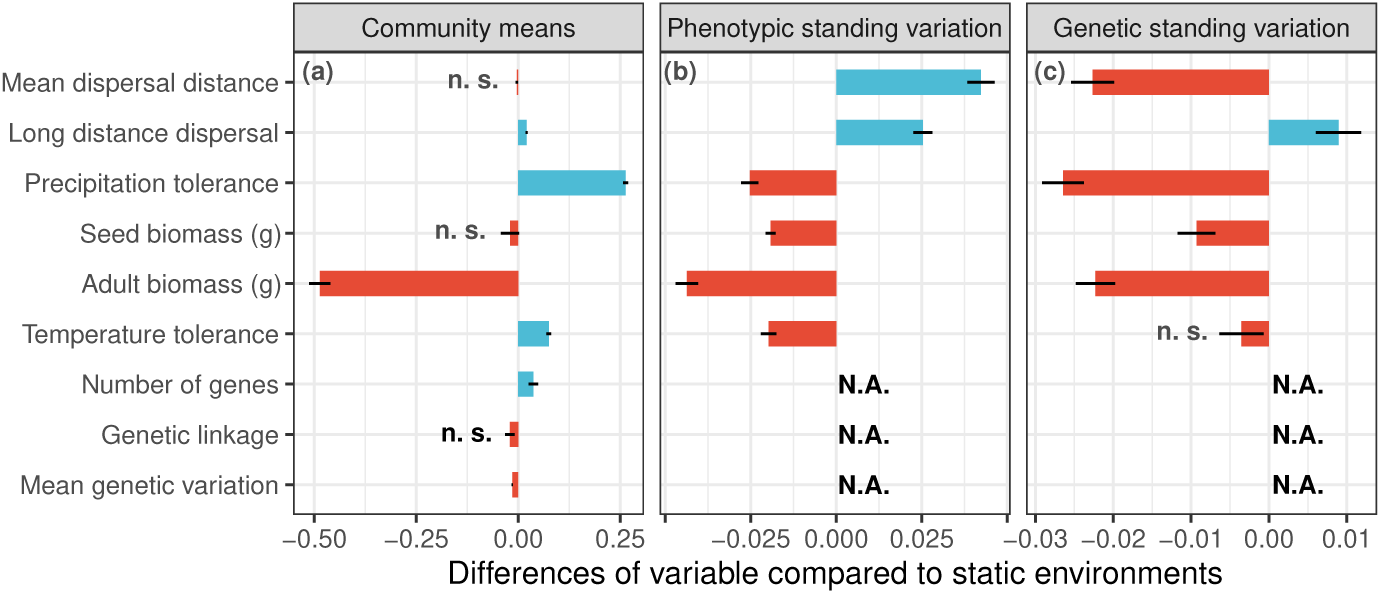
Community trait responses to temporal environmental variation along three organisational levels. (a) Differences in trait means in variable environments compared to static environments. (b) differences in phenotypic trait variances, and (c) differences in genetic trait variances in variable environments compared to static environments. Error bars show standard errors. Red and blue colors indicate negative and positive differences, respectively. Note the different axis scales. The abbreviation “n.s.” denotes differences that are not significant (*p >* 0.05). “N.A.” marks trait differences that are not available at the given level.

### Differences in standing variation (phenotypic and genetic)

Additionally to differences in the phenotypic characteristics, we found distinct patterns between environments in both phenotypic and genetic trait variation (Fig. 4(b)). While the phenotypic variation of mean dispersal distance and both phenotypic and genetic variation of long distance dispersal was increased in variable environments, all other trait variations (phenotypic and genetic) were decreased in variable environments. The trend towards a decrease in genetic variation of temperature tolerance was not significant.

## Discussion

### General differences between scenarios

Our results show how community trait composition of plant metapopulations may differ between static and temporally variable environments in a genomically-explicit eco-evolutionary model. The changing abiotic conditions in variable environments act as a constant environmental filtering mechanism (Kraft et al. 2015), where only those species survive that are able to adapt to or track environmental changes. As a result, communities are species poorer (see also Menge and Sutherland 1976). The decreased *β*-diversity furthermore suggests that these fewer species in variable environments are rather generalistic, in comparison to static environments where species seem more specialized to local environmental conditions, as evidenced by the higher *β*-diversity (cf. Gilchrist 1995). The fact that, furthermore, range-filling is reduced in the variable environments is likely a mid-domain-like effect (cf. Colwell and Lees 2000), where due to the ongoing temporal variability, the margins of a potential range will often become unsuitable quickly, impeding establishment and survival. Moreover, because the environmental change in our simulations was random rather than periodical or directional, the probability for species to find alternating suitable conditions is low. This alternating suitability, however, is the prerequisite for temporal environmental variability to favor species co-existence and increased species richness (cf. Tilman and Pacala 1993, Descamps-Julien and Gonzalez 2005). In contrast, most communities in static environments passed environmental filtering already after the first 200 years, after which species were distributed according to their environmental preferences and ecological patterns became stable.

### Study question (a): Which ecological and genomic traits enable survival in temporally variable environments?

The trait characteristics of communities in the respective environments represent successful strategies in surviving random environmental variation. The decreased values of precipitation tolerance in communities in static environments indicate increased environmental specialization. This is in contrast to communities in variable environments, where the variability in precipitation conditions favors species with higher tolerance values (i.e. specialization to local conditions are detrimental in variable environments, Gilchrist 1995, Kassen 2002). Additionally, temperature tolerance directly affects individual survival due to metabolic constraints (Fig. 2(d)). Since a high temperature tolerance decreases fitness, species are forced to keep tolerances low if they occur at their respective environmental optimum. In variable environments, this environmental optimum is hardly met. As a consequence, selection acts rather on enhancing temperature tolerance to gain long-term fitness. Therefore, our experimental design captures the evolution towards bet-hedging strategies in terms of adaptation to variable environments (Slatkin 1974).

The second aspect of survival strategies lies in the biomass patterns. In general, species in variable environments were smaller than in static environments. Since growth rates and fecundity follow MTE, smaller species are more fecund than bigger species at the expense of survival. A higher and more frequent number of offspring will spread the risk over time in variable environments (McGinley et al. 1987, Philippi and Seger 1989). Additionally, the larger range of different biomasses in static environments can be interpreted as temporal partitioning (Pronk et al. 2007), because it means that co-occurring species will reproduce at different times and intervals. This allows species to alternate dominance and thus produce temporally variable biotic conditions (cf. Olff et al. 2000, Wilson and Abrams 2005). Furthermore, both biomass and tolerance patterns suggest that specialization to avoid competitive exclusion plays a larger role in shaping communities in static environments, while communities in variable environments are primarily shaped by generalism and environmental filtering (cf. Menge and Sutherland 1976, Hulshof et al. 2013).

In order to track suitable conditions, dispersal abilities are of crucial importance in changing environments (Bourne et al. 2014). While mean dispersal distances in our simulations showed little differences between environments, long distance dispersal indeed increased in variable environments. Besides primary dispersal traits, the dispersal rate also increased in variable environments via the indirect effect of metabolic rates: the high demographic turnover that comes with higher fecundity due to decreased biomass leads to more frequent dispersal. This further explains why there was little change in mean dispersal distance between environments. With the rate of change in our simulations and the small spatial extent of our landscape, dispersal distance (which is what is controlled by dispersal traits) is less important than dispersal rate (cf. Johst et al. 2002). However, this might change in fragmented landscapes, where dispersal distance is critical to maintain connection between habitable patches (Bacles et al. 2006, Boeye et al. 2013, Bonte et al. 2010).

Lastly, species may survive by adapting to changing conditions (Jump and Peñuelas 2005). This constant adaptation requires an appropriate genetic architecture: we expected genomes to contain a high variation of trait alleles (Holt 1990) which can be recombined easily for a species to quickly respond to novel conditions (Schiffers et al. 2012, Matuszewski et al. 2015) by producing new phenotypes. Indeed, we found increased gene numbers in variable environments, which allow potentially larger range of possible expressed trait values, and thus more recombination potential. Since genetic linkage did not differ between environments, the genome size increase is due to an increased number of linkage units. Species with these larger genomes can be thought of having undergone polyploidisation or ascedent dysploidy. In fact, polyploidisation correlates with latitude and, arguably, with environmental stress (Rice et al. 2019), but direct tests of this are difficult due to feasibility (Van de Peer et al. 2017). Moreover, increased fecundity also increases adaptation potential as it leads to more recombination in a given time interval. According to our results, the adaptation response to variable environments is mainly characterised by increasing environmental tolerances. However, the changes in genomic traits did not prevent the general decrease of mean genetic variation in variable conditions, which contradicts results from a previous modeling study (Matuszewski et al. 2015). With more detailed data on the levels of variation, we will attempt to offer an explanation to this in the following section.

### Study question (b): How do temporally variable environments shape phenotypic and genetic standing variation?

Having identified survival strategies on a population phenotypic level, we wanted to know whether there are selection patterns on the standing variation within the populations — both at the phenotypic intraspecific and genetic levels. Our results enable us to identify which traits are under increased selection pressure and in which traits species benefit from variation in the different environments. Most traits, such as tolerance for environmental conditions and biomass, were more specialized, i.e., had lower variation, in variable environments at both intraspecific and genetic levels. However, it appears to be beneficial for species to maintain plasticity in dispersal distances when coping with temporal environmental variation, as evidenced by the fact that dispersal traits, especially long distance dispersal, maintained similar to higher levels of variation.

Since variation in our experiments could be increased neither by mutation (Josephs et al. 2017), nor by external gene flow (Cornetti et al. 2016), selection could only act on standing variation. Under these conditions, a higher linkage of genes preserves variation in the associated trait (cf. Teotónio et al. 2009), while low linkage genes allows faster specialization. This differentiated selection pressure might explain why we don’t see a net change in genetic linkage, because variation and specialization benefit from contrasting degrees of genetic linkage. Specialization in any given ecological function and thus the emergence of different phenotypes could also be facilitated by a low number of loci for associated traits (Schiffers et al. 2014). In contrast, phenotypic uniformity might arise from increased number of loci which stabilize variation (Fraser and Schadt 2010). Thus, the number of loci represents a potential trade-off between specialization and phenotypic robustness, which might warrant further investigation. These findings suggest that increasing the number of loci could act as a stabilizing coexistence mechanism by promoting intraspecific competition caused by phenotypic uniformity and thus greater intraspecific niche overlap. In light of this, further experiments should focus on whether phenotypic variation impacts species coexistence negatively (Hart et al. 2016) or if low species numbers first allow higher phenotypic variation (Hulshof et al. 2013).

Our results furthermore exemplify that intraspecific and genetic variation do not need to be correlated. In case of mean dispersal distance, phenotypic variation increased in variable environments. However, the genetic variation of mean dispersal distance decreased. Thus, the phenotypic variation in mean dispersal distance is due to very different phenotypes, which, in turn, exhibit relatively specialized genotypes. This further stresses the essential role of ecotypes to ensure species survival under changing environments.

### Limitations and perspectives

The fact that our simulations produced low coexistence in terms of the total number of species across the landscape might be a result of too large a trait space in the initial species pool, most of which would be filtered by the relatively narrow environmental conditions. Since the initial species pool was on average 350 species large, the probability is also high for it to contain a few strong generalist species, which outcompete other species. On the other hand, an average initial number of 10 species per grid cell means a low probability for one or more species to be sufficiently adapted to the local conditions. Nevertheless, the coexistence level obtained is also in accordance with theoretical expectations, considering that a niche partitioning along the two gradients would explain the average of four species we count in static environments (i.e. one specialized species per environmental gradient combination, see Armstrong and McGehee 1980). The filtering is also evidenced by the reduction of trait value ranges over all traits after simulation initialisation (not shown). In fact, additional post-hoc simulations with more constrained initial communities in terms of species traits resulted in a two-fold increase of surviving species numbers (not shown). This did not, however, change the general results. Small-scale disturbance or trophic interactions, e.g., herbivory could further increase coexistence, as theoretical and empirical studies suggest (Shea et al. 2004, Roxburgh et al. 2004, Chesson and Kuang 2008). But since these processes likely produce additional confounding effects, we chose not to include them in our model at this stage, albeit we identify them as potential directions for further model development. Trophic and other interactions such as mutualism, can have important effects on species survival under climate change (Berg et al. 2010) and even lead to extinction cascades if keystone species get lost (Brook et al. 2008). Since keystone species would be affected by genetic factors in the same way as any other species, our model likely underestimates net species loss effects mediated by genetic factors.

Furthermore, our model simplifies complex genetic factors and dynamics which could potentially have confounding effects on resulting patterns. For instance, linkage between genes in reality is not a binary decision, but rather a consequence of the physical distance between those genes. The larger the distance, the higher the probability of crossing over during meiosis. Additionally, genetic architecture is dynamic, especially in plants. Genomes can grow, e.g., by polyploidisation (Van de Peer et al. 2017), and shrink in size, both of which affects genetic linkage and potentially genetic variation. Since polyploidisation is often a stress response in plants it will arguably affect survival (Rice et al. 2019). Subsequent gene loss may then even initiate speciation, therefore providing new opportunities for emerging species (Albalat and Cañestro 2016). Our model hence represent the effects of genetic linkage and genome sizes without explicitly considering their respective genetic origins. Nevertheless, our findings on the interaction betweeen genetic and ecological traits call for empirical works identifying the factors that trigger these genomic processes and assessing their evolutionary relevance (Van de Peer et al. 2017).

To make our model and the findings on genomic and ecological traits under temporal evironmental variation more applicable and relevant to real-world systems, the model could be constrained by real data in further studies. For instance, the model could be initiated with simulation arenas which can be directly derived from actual landscapes, including environmental conditions (e.g. from Karger et al. 2017). Species-specific parameters could be taken from databases for phenotypic traits (Kattge et al. 2011) and occurence records (GBI) and enriched by genomic information (Dong et al. 2004, Howe et al. 2020) to constrain initial parameter space for the creation of random communities. Thus, our model represents an oppurtunity to integrate different datasets from genomes over traits and occurrences to environmental in a single mechanistic framework.

Even in the current state, our model addresses a number of eco-evo-environmental phenomena (cf. Govaert et al. 2019). The emerging patterns additionally inspire new hypotheses which can be used to guide fieldwork and experimental studies. The consideration of genomic traits, for example, implicates the explicit consideration of new perspectives on biodiversity dynamics during impending climate change (Fig. 5). For scenarios of short-term change of environmental conditions, i.e., warming, lower or increased precipitation and more frequent extreme events, adaptation can only exploit standing intraspecific or genetic variation, rather than novel mutations. Species with high phenotypic variation will likely have good adaptation potential, regardless of genetic characteristics. For species with low phenotypic variation, adaptation potential depends on genomic traits. Species that have highly specialized, i.e., uniform, phenotypes, and show little or no genetic variation will only be able to survive rapidly changing conditions by tracking their specific favourable conditions. Fragmented environments or poor dispersal abilities therefore will likely lead to the extinction of those species. Even if species have high genetic variation, genetic architecture is crucial for their performance. With a high degree of genetic linkage, species might not be able to adapt critical traits in time to react to changing conditions, since a beneficial trait allele might likely be linked to other disadvantageous trait alleles. Thus, net fitness is unlikely to increase. Low linkage, on the other hand, might lead to species who quickly adapt to new environments as they are not impeded by genetic hitchhiking. However, if linkage is too low, species will also quickly lose genetic variation, rendering them unfit to react to subsequent change. Any conservation measures targeted at particular species should thus consider population structure and genomic traits of species. Hence, while the importance of genetic diversity is already acknowledged in conservation biology (Ramanatha Rao and Hodgkin 2002) — additional to functional diversity (Diáz and Cabido 2001), it is genetic architecture that will determine adaptation success.

**Figure 5.**
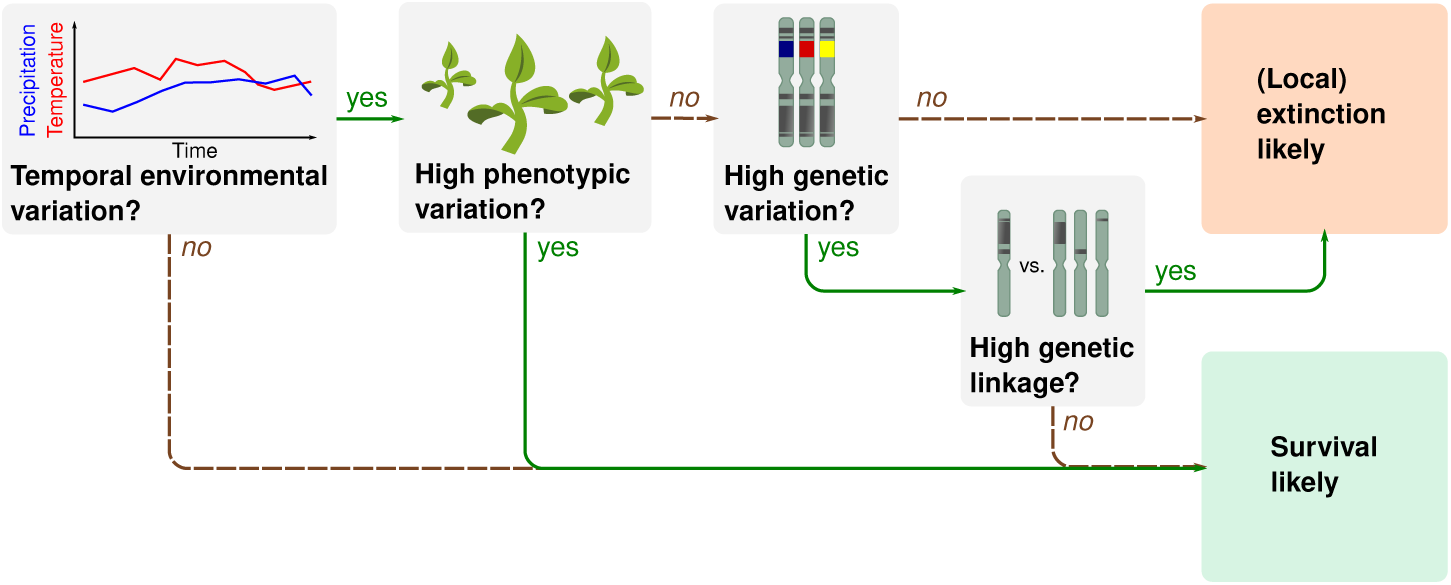
Ecological and genomic factors influencing species survival under variable environmental conditions.

## Conclusion

In this study we have demonstrated complex interactions between genetic and ecological realms by using a simulation model that explicitly considers genetic architecture of plant communities in changing environments. These eco-evolutionary feedbacks broaden our understanding of the role of trait-specific standing variation in species survival and adaptation (Fig. 5). This enabled identifying ecological strategies of species to survive variable environmental conditions. Variable environments select species with higher tolerances and faster life cycles while species maintain variable dispersal abilities that facilitate tracking favorable environmental conditions. These adaptations are, however, mostly enabled by large genomes, which allow maintaining a high degree of genetic variation. Furthermore, we could show that selection pressure differs between traits and that there might even be positive selection pressure to maintain higher genetic variation for dispersal traits.

Our findings suggest that genomes are subject to opposing forces — especially under changing conditions. While constant environmental filtering impoverishes genomes, there is a selective force to maintain variation in the genome to adapt for future change. This conflict can be mediated to a certain degree by genetic architecture, namely a higher number of genes which allows more genetic variation and a high linkage of loci which impedes the loss of variation. However, traits that need quick specialization require a low number of weakly linked loci. These complex interdependencies of genomic traits may thus further promote the high diversity in genetic architecture and ecological strategies in real-world species.

Additionally, our theoretical approach provided potential mechanisms responsible for the incongruence of phenotypic and genetic variation, which is sometimes found in nature. A mechanistic link between negative correlation in those types of variation means that special care is called for when inferring genetic variation from phenotypic variation and vice-versa.

In summary, this study highlights the importance of genomic traits for the functional assessment of local populations, species and metacommunities. We hope that conservation studies make more use of these characteristics to prioritize conservation efforts and expect future studies to investigate the genetic architecture of specific traits in natural populations.

## Acknowledgements

We thank Daniel Vedder for invaluable edits and additions to the model’s code and for compiling the model’s documentation. We thank Ludmilla Figueiredo, Thomas Hovestadt, Sonia Kéfi, Anne Lewerentz and Daniel Vedder for helpful comments on previous versions of this manuscript.

## Contributions

JSC designed the research with input by LL. LL implemented the model, ran the simulations, performed the analysis and wrote the manuscript. JSC contributed to the writing of the manuscript.

## Code availability

The model code, experiment definition files, and analysis scripts are available at https://github.com/lleiding/gemm.

## Data availability

The results should be reproducible by using provided simulation codes and configuration files (see Code availability). Any data and codes associated with this study will be made accessible upon request.

